# Cytosolic Enhanced Dark Epac-Based FRET Sensors for Intracellular cAMP detection in live cells via FLIM

**DOI:** 10.1101/2024.09.30.615785

**Authors:** Giulia Zanetti, Jeffrey B. Klarenbeek, Kees Jalink

**Affiliations:** Division of Cell Biology, The Netherlands Cancer Institute, Amsterdam, The Netherlands; van Leeuwenhoek Centre of Advanced Microscopy, Amsterdam, The Netherlands; Swammerdam Institute of Life Sciences, University of Amsterdam, Amsterdam

**Keywords:** FLIM, FRET, EPAC, cAMP, biosensors, live cells

## Abstract

Fluorescence Resonance Energy Transfer (FRET)-based biosensors are powerful tools for studying second messengers with high temporal and spatial resolution. FRET is commonly detected by ratio-imaging, but Fluorescence Lifetime Imaging Microscopy (FLIM), which measures the donor fluorophore’s lifetime, offers a robust and more quantitative alternative. We have introduced and optimized four generations of FRET sensors for cAMP, based on the effector molecule Epac1, including variants for either ratio-imaging or FLIM detection. Recently, Massengill and colleagues introduced additional mutations that improve cytosolic localization in these sensors, focusing on constructs optimized for ratio-imaging. Here we present and briefly characterize these mutations in our dedicated FLIM sensors, finding they enhance cytosolic localization while maintaining performance comparable to original constructs.

## Introduction

Second messengers are a heterogeneous group of molecules involved in signal transduction and intracellular regulation in response to various stimuli. The first identified second messenger, cyclic Adenosine Monophosphate (cAMP), interacts with several downstream effectors such as protein kinases, ion channels and Guanosine TriPhosphate (GTP) exchangers to control a large variety of biological processes, including gene transcription, cell metabolism, proliferation, development, as well as more specialized functions depending on the cell type [1,2]. cAMP is synthesized from Adenosine TriPhospate (ATP) by members of the Adenylate Cyclase (AC) protein family, which in turn are directly controlled by G Protein-Coupled Receptors (GPCRs) through stimulatory (Gα_S_) and inhibitory (Gα_I_) G proteins. Conversely, the breakdown of cAMP is through a family of cyclic nucleotide PhosphoDiesterases (PDEs) [1-4]. Production and degradation of cAMP are tightly regulated, resulting in highly dynamic and agonist-specific alterations in cAMP concentration within the cell. Additionally, spatial distribution of cAMP within cells reveals a non-homogeneous compartmentalization, with cAMP concentrated near the plasma membrane, nucleus and mitochondria, essential for ensuring efficient and precisely localized cellular responses [5,6].

FRET-based biosensors have emerged as excellent tools to study second messengers with high temporal and spatial resolution [7-9]. FRET, i.e. the non-radiative transfer of energy from an excited donor (D) fluorophore to a suitable acceptor (A), is strongly distance-dependent in the range of 1–10 nm. FRET changes are commonly detected by ratio-imaging of D and A intensities. However, ratiometric readouts are not fully quantitative because the ratio is affected by factors such as fluorophore concentrations, background fluorescence, photobleaching, light source fluctuations as well as by characteristics of the optical filters used to isolate D and A fluorescence [7,9]. Therefore FLIM, an alternative and much more quantitative manner to detect FRET [3, 9-13], is rapidly gaining interest. FLIM focuses only on an intrinsic property of the donor fluorophore’s lifetime, i.e. the time that it spends in the excited state before returning to the ground energetic level by emitting a photon [13,14]. As the excited state lifetime of the donor is linearly related to the FRET efficiency, FLIM provides more accurate and reliable FRET measurements regardless of the limitations mentioned previously. With commercial FLIM instrumentation now widely available, it also becomes essential to design FRET sensors specifically tailored for lifetime detection.

We and others [3, 15-20] originally described FRET biosensors for ratio-detection in which the cAMP effector protein Epac1 (Exchange protein directly activated by cAMP) is sandwiched between a cyan donor and yellow acceptor fluorescent protein. Binding of cAMP to this sensor partially unfolds Epac1, leading to an increased distance between D and A, and therefore, a decrease in FRET. Our design was based on full-length Epac1 because we found this to display largest FRET changes, and we introduced mutations that render the Epac1 moiety catalytically dead and that prevents its localization to the plasma membrane, delta-DEP (Disheveled, Egl-10 and Pleckstrin) [19,21]. In subsequent studies, we optimized choices for D and A fluorescent moieties as well as linkers, and we addressed D-A orientations, fluorophore maturation as well as phase separation issues [18]. We also introduced sensors with highly extended dynamic range and the first sensors specifically optimized for detection by FLIM [19].

Our fourth-generation Epac sensors, which include higher affinity versions, have unparalleled FRET contrast and are widespread in use in the field. For example, the EPAC sensor H187 (construct termed Epac-S^H187^) contains a Q270E mutation to boost its affinity to ∼ 4 µM and it includes a tandem of the circular permutated YFP variant, ^cp173^Venus, as acceptor cassette and the brightest and most bleaching-resistant CFP variant, mTurquoise2, as a donor. A close relative, our Epac-S^H189^, has a Y145W mutation in both Venus moieties which renders the acceptor dark, thus allowing for optimal detection by FLIM, which makes this the construct of choice for co-expression with additional FRET/FLIM sensors in the same cells [20] (Fig. 1A).

**Fig 1.**
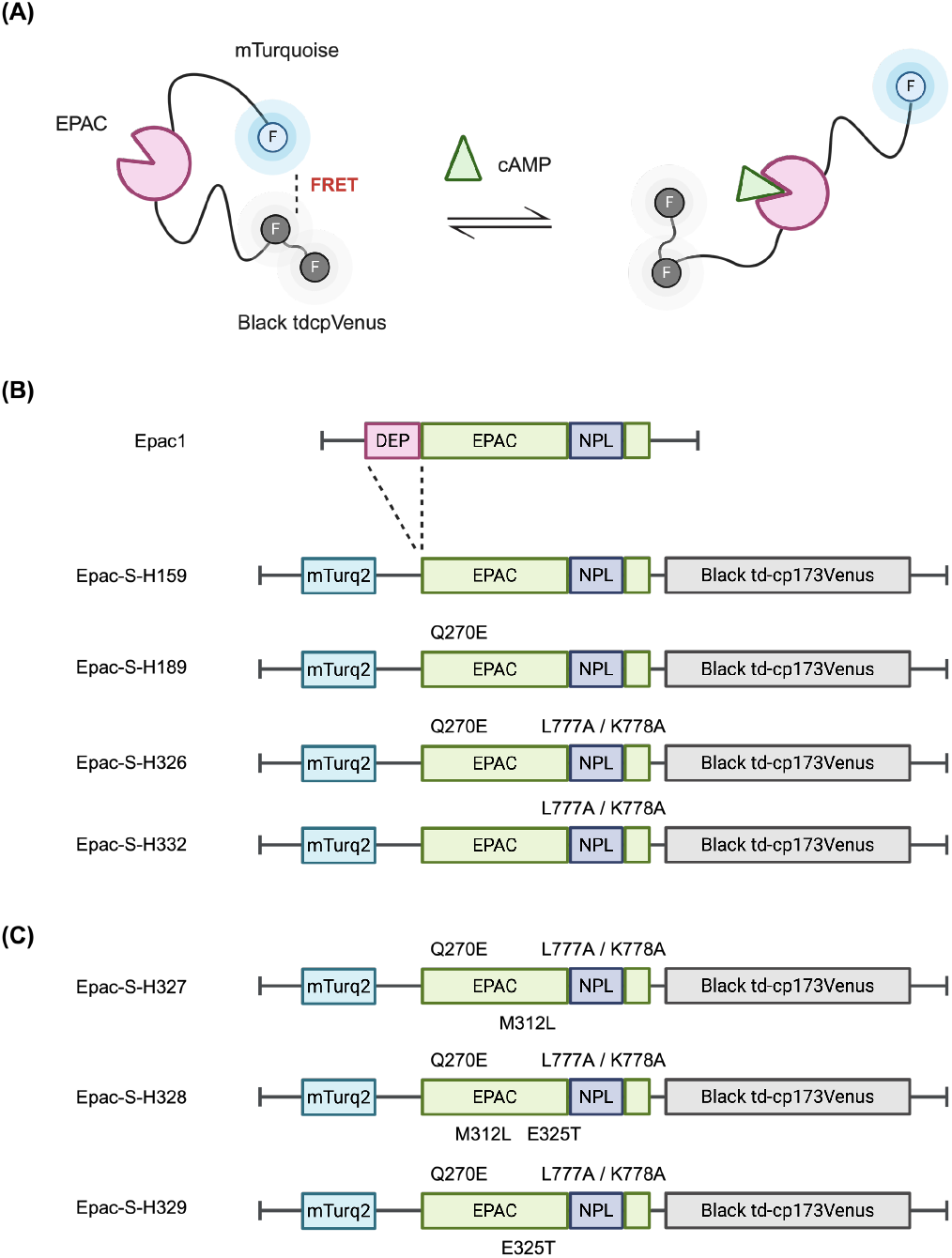
Design and mutational overview of the Epac constructs. (**A**) Schematic representation of the FRET-FLIM Epac sensor, dark-acceptor variant, illustrating its main structural domain and functional features. **B** and **C** present a series of sensor constructs, each associated with a linear plasmid map detailing the specific mutations introduced relative to the Epac1 sequence. The maps highlighting the position of the mutations, providing a comprehensive overview of the genetic modifications applied to generate the sensor variants.

All of these sensors are predominantly localized in the cytosol, but they also bind to the nuclear envelope to a small degree. Whereas this is prominent only in low-expressing cells, it might affect the response to cAMP in unpredictable ways. Starting from our Epac-S^H187^, Massengill and colleagues identified the residues responsible for localization to the nuclear envelop and introduced mutations (L777A/K778A) that render the sensors 100% cytosolic, which they renamed to cAMPFIREs. They also introduced mutations to further increase the affinity for cAMP, i.e. M312L and E325T, which is crucial for application in cells that exhibit low baseline cAMP levels and small agonist-induced changes in cAMP concentration, such as neuronal cells. The numbering of the mutations reported here refers to the Epac1 moiety structure; however, these mutations correspond to different positions in the sensors’ sequences (Tab. 1). For consistency and ease of reference, the original numbering will be maintained throughout this report.

In this brief report we describe introduction of those useful mutations in our dedicated FLIM sensors, starting from Epac-S^H159^ and Epac-S^H189^ (Fig. 1B). As expected [17], we find that these sensors localize cytosolically even in low-expressing cells, and we find that they perform as well as the original sensors in all aspects tested. For completeness we also introduce dedicated FLIM variants with the highest-affinity sensors with M312L and E325T mutations (Fig. 1C). All new variants will be deposited at Addgene.

**Table 1.** Summary of the mutations mentioned in this study and their corresponding positions within the sensor constructs. The left column displays the mutations based on the Epac1 moiety structure, while the right column indicates the respective positions in the sequences of the sensor variants. Although the numbering of mutations is originally referenced to the Epac1 moiety, for consistency and ease of reference, this numbering is maintained throughout the report to facilitate comparison between the Epac1 structure and the sensor sequences.

| Mutations |  |
| --- | --- |
| Epac1 Moiety WT | Epac Sensor |
| Q270E | Q357E |
| E325T | E412T |
| M312L | M399L |
| L777A | L864A |
| K778A | K865A |

## Results and Discussion

### Dedicated FLIM sensors with improved cytosolic localization

We focused on our original design FLIM sensors with normal (9.5 µM; Epac-S^H159^) and high (4 µM; Epac-S^H189^) cAMP affinity, because these affinities are most relevant for experiments in non-neuronal cells. While basal and stimulated cAMP levels in some neuronal cells are so low that they necessitate higher-affinity sensors, in the cultured cells that are widely in use in cell biology and oncology research, basal cAMP concentration is in the high nanomolar range and stimulated levels may reach up to tens of µM. Thus, in these cells H159 and H189 are predicted to provide the largest FLIM span and to most reliably report high cAMP levels. Based on Epac-S^H159^ and Epac-S^H189^, we prepared constructs harboring both the L777A and K778A mutations using a linear insertion and ligation cloning strategy (see methods) yielding the new sensors Epac-S^H332^ and Epac-S^H326^ (Fig. 1B), respectively; the performance of these variants will be described in this report. For completeness, several additional mutants were also generated (Fig. 1C), however, these variants are out of the main scope of this brief report.

Epac-S^H332^ and Epac-S^H326^ were compared to the parental constructs Epac-S^H159^ and Epac-S^H189^ by transient expression in HEK 293T cells. As reported, introduction of the Massengill mutations L777A and K778A completely abolished all detectable binding to the nuclear envelop (compare Fig. 2A and B to Fig. 2C and D). The same observation was made when sensors were expressed transiently in Neuro2A, Cos7, A431, and HeLa cells (*see supplemental information, Fig. S1*).

**Fig 2.**
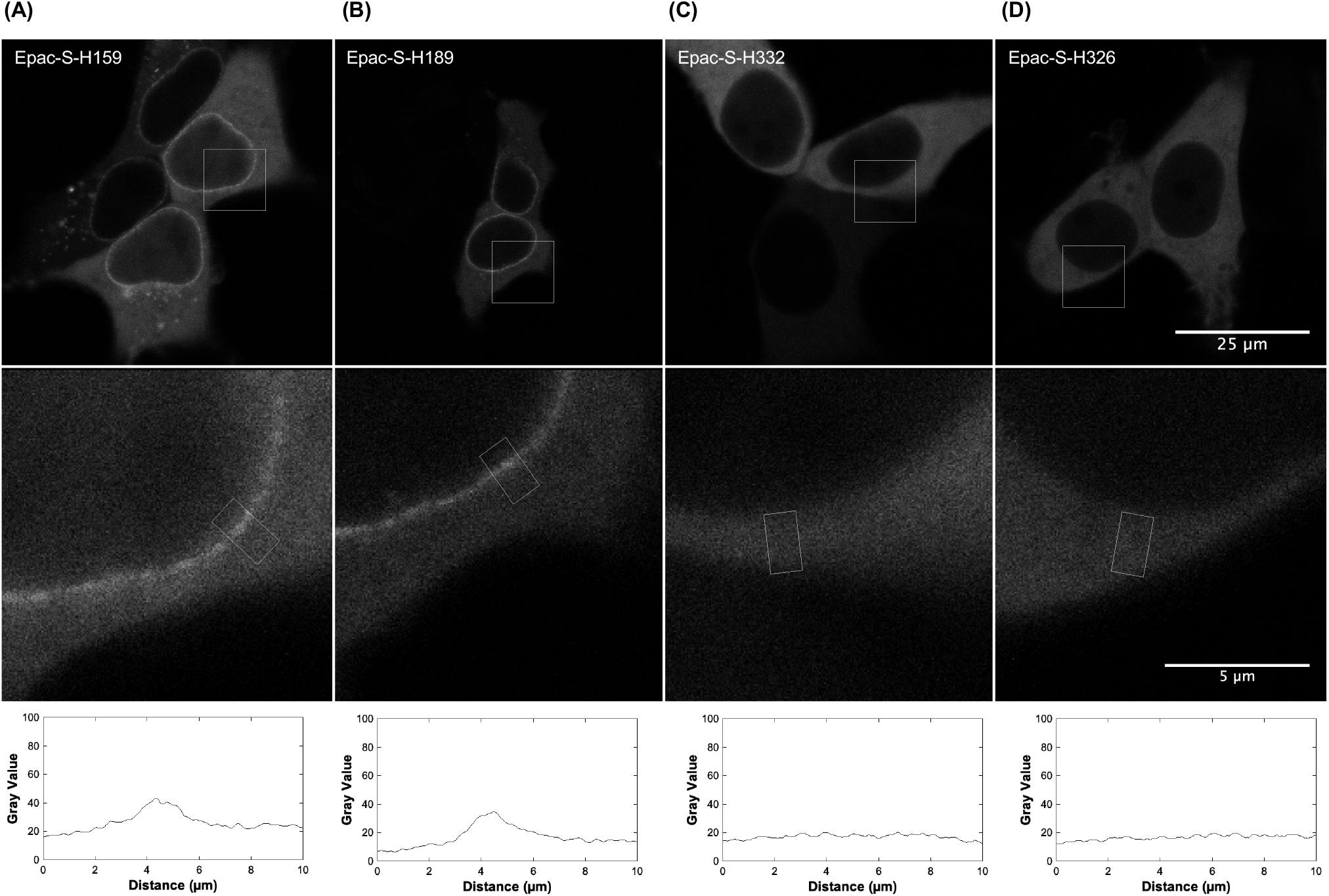
Representative microscopy images displaying the subcellular localization of Epac sensors within HEK cells. The parental sensors (**A**-**B**), Epac-S^H159^ and Epac-S^H189^, show a clear enrichment at the nuclear membrane, whereas the mutated variants (**C**-**D**), Epac-S^H332^ and Epac-S^H326^, lack this envelope association. Below each image, magnified views of the regions marked by white squares are shown. In the bottom panel, distance (µm) versus gray value plots are presented, derived from intensity analysis performed in Fiji on the white rectangular regions in the direction perpendicular to the nuclear envelop. Images were taken with a 63x, 1.4 N.A. oil immersion objective.

Massengill and colleagues also reported a susceptibility of the ratiometric variants of our high-affinity sensors, Epac-S^H187^, to form aggregates in the cytosol. These authors showed ‘typical’ images of cells with large fluorescent clusters with aberrant FRET efficiency and lifetime [17]. This is to our surprise because in thousands of experiments over the years, carried out in a variety of cultured cell lines including HEK 293T cells we never observed a noticeable problem with large aggregate formation. In fact, the design with a double Venus acceptor moiety was introduced specifically to abolish formation of bright fluorescent speckles in the earlier variants [20]. Aggregate formation was also not mentioned in the feedback from hundreds of users of our sensors. Nevertheless, we systematically compared localization in hundreds of cells expressing the dark-acceptor FLIM sensors to assess this issue. Overall, no significant difference in sensor aggregation was detected (Fig. 3A, B) except, perhaps, in some cells with unpractically low expression of the sensors. Thus, at least in our hands and our cells, we find no evidence that the L777A/K778A mutations are an important determinant of aggregate formation.

**Fig 3.**
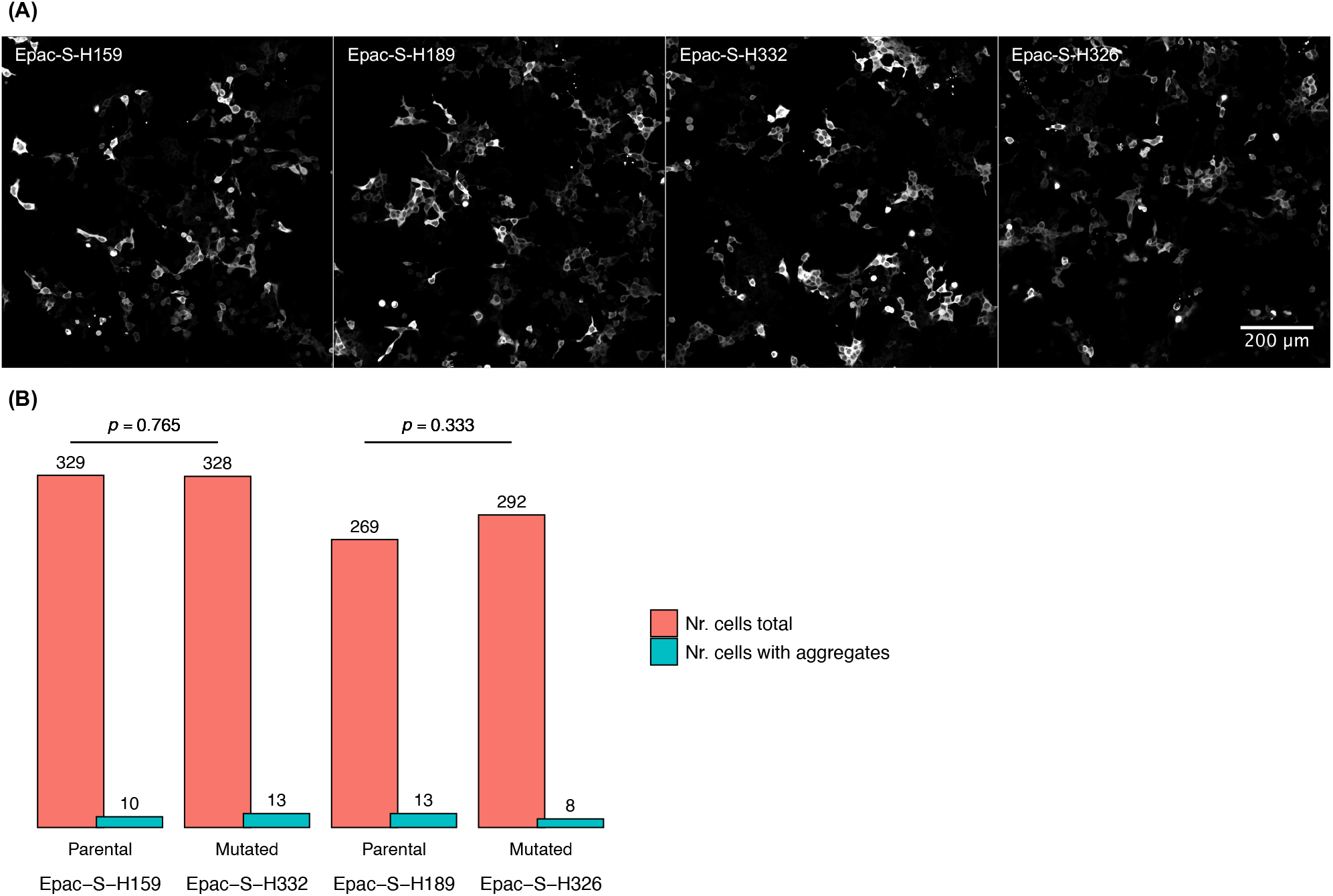
Analysis of aggregate formation in parental and mutant Epac sensors. (**A**) Fluorescence microscopy images acquired with a 63x, 1.4 N.A. oil immersion objective, 4×4 field mosaic, providing an overview of HEK cells lines expressing Epac-S159, Epac-S-H189, Epac-S-H332, and Epac-S-H326. A comparison of the images using Fiji, along with the quantification of total cells and cells with aggregates (**B**), demonstrate that the introduced mutations (L777A/K778A) in the Epac sensors do not significantly affect aggregate formation. Aggregates formation was scored blindly by zooming in to the images and eye-balled assessment of aggregates. Binomial tests indicate no significant difference in the probability of aggregate formation between parental (H159, H189) and mutant (H326, H332) sensors (p >> 0.05).

### Evaluating the performance in FLIM experiments

Mutations in the Epac backbone may also affect the protein conformation, and thereby, the FRET efficiency of the sensors. We therefore systematically characterized the FLIM response of our novel set of dedicated dark-acceptor sensors, again by comparing them to the unmutated parental constructs. Performance was evaluated by comparing agonist-induced changes in transiently transfected HEK 293T cells stimulated with isoproterenol (ISO), a synthetic catecholamine that acts as a beta-adrenergic agonist leading to an increase in intracellular cAMP levels. In the same cells we also investigated the response span of the FLIM sensors by comparing baseline lifetime to maximal lifetimes obtained after addition of forskolin (FSK), which directly activates AC, and 3-isobutyl-1-methylxanthine (IBMX) to block the PDE that break down cAMP. For these experiments, time-lapse images of large numbers of cells were collected. Cells were segmented and average lifetime of individual cells were analyzed essentially as published [13]. Figure 4A shows the lifetime traces of each cell as well as box plots that depict the average baseline (first 100 seconds) and maximal lifetimes (last 100 seconds) of these cells.

**Fig 4.**
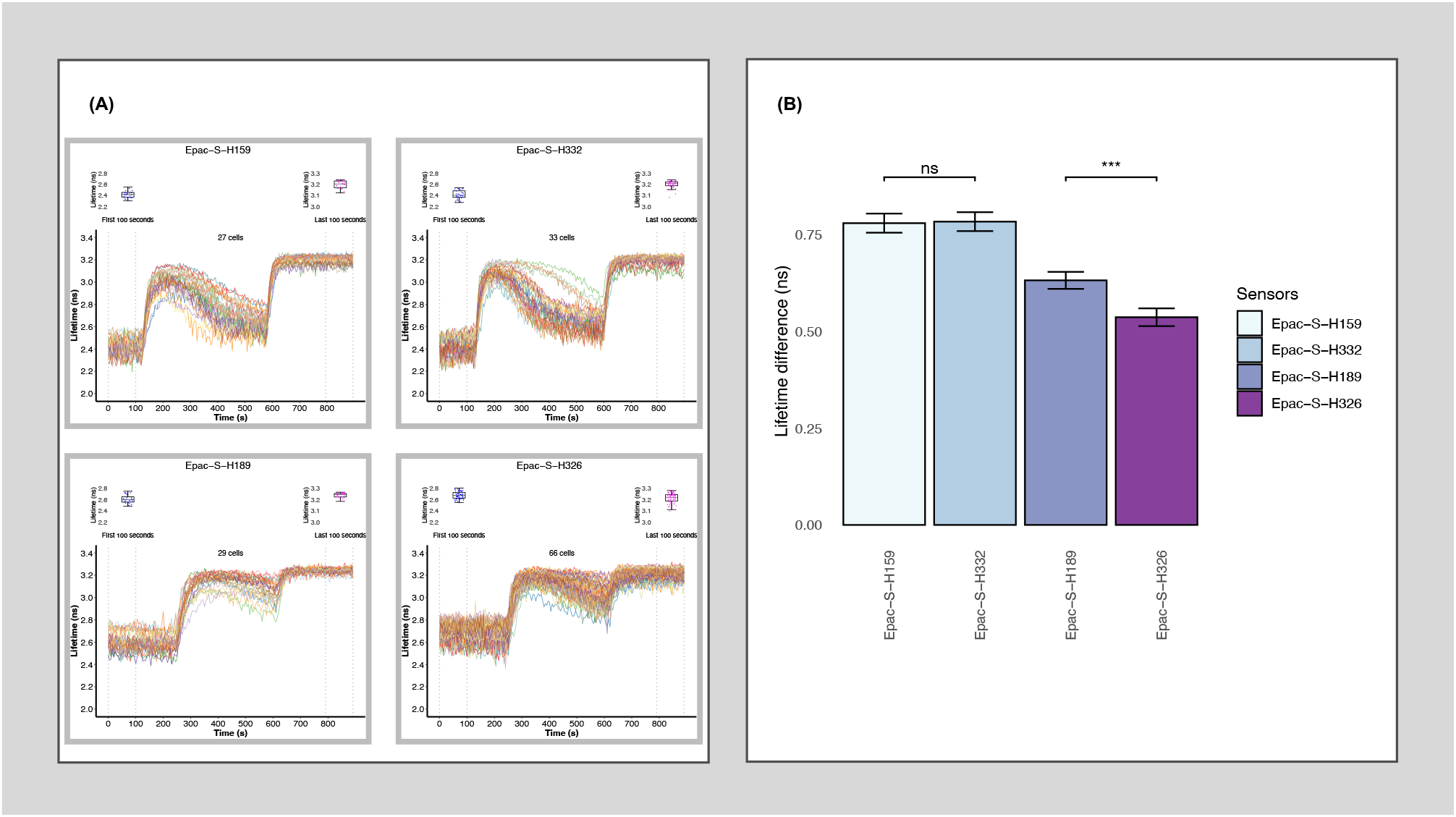
Detection of cAMP dynamics in HEK cells by FLIM. (**A**) Single cells were segmented using the Cellpose algorithm and FLIM time-lapse traces are shown from cells expressing the FRET-FLIM sensors Epac-S^H159^, Epac-S^H332^, Epac-S^H189^, and Epac-S^H326^. Cells were imaged at rest and following stimulation with ISO (100 nM) added at 100 seconds for the normal affinity sensors and at 250 seconds for the high affinity sensors, and FSK (25 μM) + IBMX (100 μM) added at 600 seconds. The traces illustrate dynamic changes in fluorescence lifetime, with corresponding box plots highlighting both the basal and stimulated states of cAMP dynamics in live cells. 20x, 0.75 N.A. dry objective. (**B**) Bar plot illustrating the maximum FLIM span reached by each Epac-S variant expressed in HEK cells (p = 1.924 x 10^-6^, Wilcoxon test).

Whereas the L777A/K778A mutations clearly abolished the partial localization of sensors to the nuclear envelop, they did not detectably alter FLIM response kinetics or response amplitude upon receptor stimulation. It is not straight-forward to detect sensor response kinetics from these experiments. In the majority of cells, the lifetime response peaked within 50 s after addition of ISO, but these kinetics predominantly reflect the time required to activate the receptor and signaling cascade. We therefor conducted experiments in which the response to rapid release of cAMP from a caged precursor, DMNB (4,5-Dimethoxy-2-Nitrobenzyl)-caged cAMP, was recorded. These experiments showed that mutant as well as parental constructs bind to cAMP within 0.2 s (Fig. 5). Importantly, no differences in response rate (K_on_) were observed in constructs harboring the L777A/K778A mutations.

**Fig 5.**
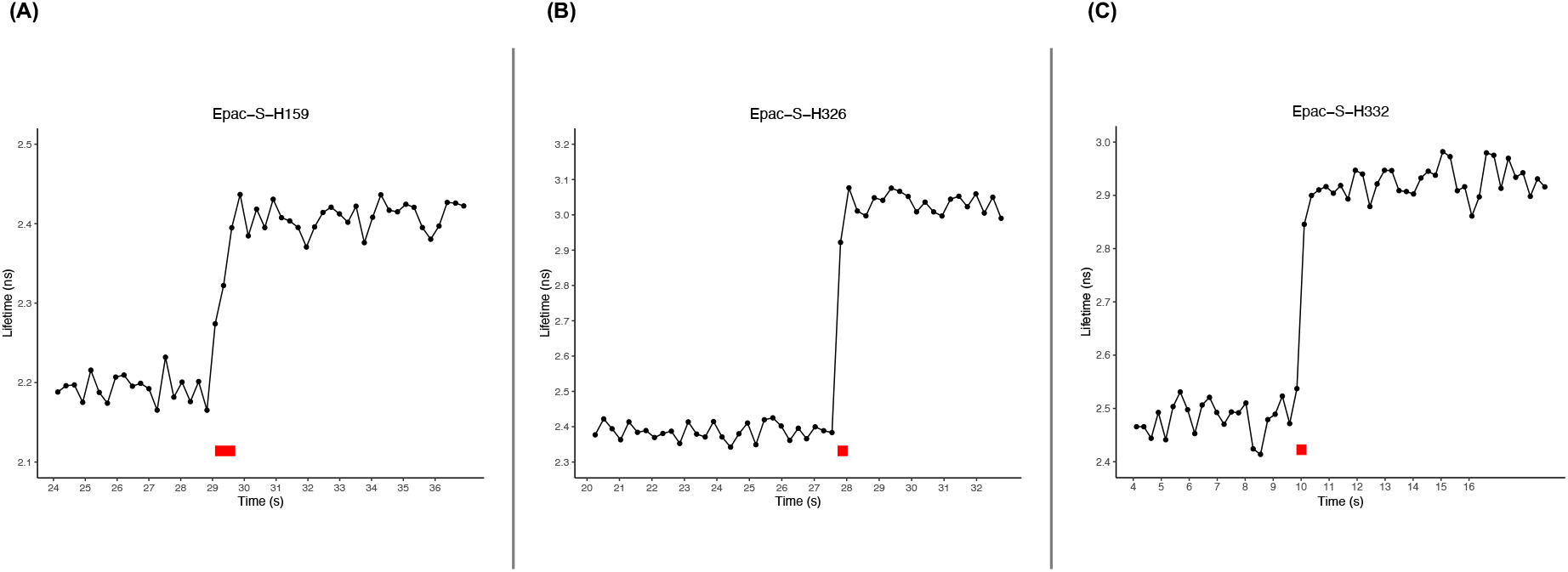
Response of selected Epac sensors to flash photolysis of caged cAMP. Responses are shown for (**A**) H159, (**B**) H326 and (**C**) H332. After recording of a baseline, a flash of UV light was applied by manually operating the shutter as fast as possible (red bars). As the duration of the flashes was quite poorly reproducible, shown examples are selected responses showing the maximum response rate of individual cells.

As stated, the L777A/K778A mutations may affect backbone folding and thereby the FRET efficiency (E). Because E relates linearly to lifetime τas: 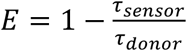 we quantitatively compared the sensor lifetime before and after stimulation with IBMX and FSK. Basal and stimulated lifetimes of the mutated normal affinity sensor, Epac-S^H332^ did not significantly differ from its parental construct Epac-S^H159^, and consequently the FRET span was identical too. In contrast, the mutated version of the high affinity construct Epac-S^H326^ differed displayed a slightly higher baseline lifetime than its parental construct Epac-S^H189^ in HEK cells, likely due to its somewhat higher affinity [17] (Fig. 4A, B). Because lifetimes after stimulation with FSK and IBMX did not differ, H326 displayed a smaller FLIM span than H189 in these cells. Note however that this is due to the combination of high baseline cAMP levels in HEK cells and relatively high sensor affinity. A similar observation was made in HeLa cells, Cos7 cells and Neuro2A cells. We conclude that the Massengill mutations reliably remedied localization to the nuclear envelop while not degrading the FRET span of the sensors appreciably.

Massengill and colleagues also presented 2 mutations that further increase the affinity of our high-affinity construct, namely M312L and E325T. Whereas such high affinity variants are of little use in our cells due to normal baseline cAMP concentration, we have also prepared dedicated FLIM sensors that include M312L, E325T and both together (Fig. 1C). The resulting constructs, Epac-S^H327^, Epac-S^H328^ and Epac-S^H329^, appeared functional when expressed in cells and were purely cytosolic (*see supplemental information, Fig. S2, S3*). These constructs will also be deposited at Addgene for use in cells with very low cAMP levels such as neuronal cells, but note that our cells do not permit proper characterization of their dynamic behavior or their FLIM span. We also note that the constructs H327 and H329 are just dark-acceptor variants of cAMPFIRE-M and H.

### Concluding remarks

We present and briefly characterize a new set of dedicated FLIM sensors which include the L777A/K778A mutations introduced by Massengill and colleagues to diminish binding to the nuclear envelop of the sensors. We partly characterize the new constructs, focusing on aspects expected to be affected by the aforementioned mutations, i.e., the localization within the cells and the FRET/FLIM characteristics: FRET E at resting cAMP levels, FRET span (maximum lifetime difference after full activation of the cells) and response kinetics. We did not focus on characteristics pertaining to the fluorescent protein (FP) moieties, such as bleaching and maturation, nor did we address specificity for cAMP and other characteristics which were reportedly unaltered by these mutations [17].

As expected, the new sensors H326 and H332 display a homogeneous distribution throughout the cytosol. Massengill et al report that the L777A/K778A mutations correct for two separate flaws: sensor localization to the nuclear envelop, and a tendency to form aggregates/clusters throughout the cytosol. As mentioned in Results, we did never find evidence for sensor clustering upon transient or stable expression of the sensors, perhaps due to selection of higher expression levels as compared to Massengill et al. However, if the speckles they observe in dim cells are in fact protein aggregates, one would expect this to become worse with increasing cytosolic concentration of the sensor. The opposite is observed. We speculate that cytosolic speckles do not represent sensor aggregation, but that they may in fact reflect saturable binding to membranes which contain the same (or similar) binding partners as those present on the nuclear envelop. Such membranes may be separate vesicles or perhaps contiguous with the nuclear envelop, like the endoplasmic reticulum (ER). This is in accordance with observations from Parnell and colleagues [22] for wildtype Epac1, although these authors do not comment on it. The exact nature (i.e., what proteins are involved) of the binding of Epac1 to the nuclear envelop is unresolved and further analysis is outside the scope of this brief paper.

Our lifetime analysis revealed similar performance between the new sensors and their parental constructs, with a slightly higher baseline value (2.7 +/-0.1 ns) for the high-affinity mutated sensor as compared to the parental construct (2.6 +/-0.1 ns). This is in accordance with the slight increase in affinity introduced by the L777A/K778A mutations reported by Massengill. All sensors reached 3.2 ns upon calibration with FSK and IBMX, and thus presented a robust FLIM span. These data are also in line with our observation that the sensors which specifically decorate the nuclear membrane do not display a different lifetime. In all, we recommend the new FLIM sensors for future research due to their even cellular distribution, the robust and quantitative readout by FLIM, and the increased options for co-expression with other FRET or dye sensors caused by to absence of bright acceptor emission in the YFP channel.

Note that in the present study we found no evidence for the claim of ‘improved sensitivity’ laid out in Massengill et al. This is because the high basal concentration of cAMP in most cultured cell lines already partly (or largely) saturates the cAMPFIRE-M and H sensors and stimulation with agonists such as isoproterenol, even at 1 – 10 nM, often fully saturates them, compromising the analysis of response kinetics. Sensitivity refers to the ability to detect minor deviations in the analyte, which requires selection of a sensor with the proper Kd to detect such changes. Thus, analysis of cAMP changes in our cells requires the normal or (our) high-affinity (= cAMPFIRE-L) sensors, and application of the cAMPFIRE-M and H saturated the response to the point that it was impossible to determine a proper FLIM span *in vivo*. Yet, as a service to the community we prepared and briefly tested the FLIM-optimized cAMPFIRE-M and H constructs and made them available via Addgene. Once again, note that we do not have the cells to properly test their FLIM span.

One major difference between this study and that by Massengill is that we used FLIM detection throughout our analysis. FLIM readout readily allows us to quantitate FRET efficiency, and, using a one-time calibration curve, translate that to cAMP concentration. This quantification is than valid for all FLIM setups throughout the world. In contrast, ratio imaging is a pseudo-quantitative technique. Proper calibration can make ratio-imaging quantitative for one particular setup, but microscope shading effects, filter ageing and chromatic aberrations still need to be accounted for [23]. Ratio imaging is commonly normalized by setting the baseline (resting) ratio to 1, ignoring individual variations in cell resting cAMP concentration and adding unnecessary variability. Also in this respect, FLIM analysis is much more straight forward.

On top of that, our study appears to rely on cells that express intermediate levels of the sensors as opposed to the low expression levels (as judged from the fact that we only observe nuclear envelop localization in the dimmest cells) used by Massengill and colleagues. While high expression of sensors is more likely to affect cAMP kinetics (buffering), low-intensity cells yield much more noisy traces and will increase variability due to the presence of cellular autofluorescence, both in ratiometric and in FLIM detection.

A final caveat concerns the localized nature of cAMP signals. It has by now been well established that cAMP signaling is highly compartmentalized in cells and that very local and very high cAMP concentration may occur in small nanodomains [24-26]. The analysis of average cAMP concentration throughout the cells in studies like the present and that by Massengill et al completely ignores the nuances of spatiotemporally localized cAMP signals. Individual microdomains may in fact require sensors with different affinity to properly detect dynamics. On top of that, cytosolic sensors are more likely to disrupt localization of signaling because they protect diffusing cAMP molecules against the action of PDEs which are thought to be pivotal in maintaining cAMP gradients. Thus, to obtain a more comprehensive view of cAMP signaling likely necessitates the use of affinity-tailored and locally anchored sensors, rather than cytosolic versions.

## Materials and Methods

### Cell culture and transfections

HEK 293T cells were cultured in DMEM (Gibco, 31966-021), supplemented with 10% FCS (Gibco, 10270) and 1% Penicillin/Streptomycin (Sigma-Aldrich, P4458), at 37°C in a humidified incubator with 5% CO_2_.

24 hours before imaging, cells were seeded on 24-mm diameter coverslips in 6-well plates at approximately 20% and transient transfected with 1 µg DNA per well using FuGene6 Transfection Reagent (Promega, E2692).

### Cloning

A novel set of FLIM sensors were created using either side directed mutagenesis (SDM) or insertion of a gene block. Starting from our parental Epac-S^H159^ and Epac-S^H189^ plasmids, we introduced the nuclear delocalization mutations L777A/K778A by SDM. The following primers were used: Fw TTCATGCCCCTTCTTgcCgcAGACATGGCCGCCATT and Rv AATGGCGGCCATGTCTgcGgcAAGAAGGGGCATGAA. Additionally, for the super high affinity sensors, the M312L and E325T mutations were introduced via the insertion of gene blocks. All mutations were checked by sequencing using primers CACGACTGGAGCCTCTTCAACA and TTCGAAAGCCCCCAGGTCA for L777A/K778A and M312L or E325T respectively. All constructs and their sequences have been deposited at Addgene.

### Confocal microscopy and Fluorescence Lifetime Imaging

Experiments were performed in 2 mL HEPES buffered FluoroBrite DMEM (Gibco, A18967-01), kept under 5% CO_2_, at 37°C in a stagetop incubator (Okolab, H301-K-FRAME). Here we used Leica Stellaris 8 FALCON confocal FLIM microscope using LAS X software, equipped with fast time correlated single photon counting detector(s). For FRET-donor excitation we used the 448 nm line of the white light laser, pulsed at 40 MHz. The spectral donor emission was collected using the Power HyDX2 detector, in the spectral range between 460 – 540 nm.

Imaging to evaluate the sensors’ localization and aggregation tendency was performed using a 63x, 1.4 N.A. oil immersion objective in xyz-mode. Xyt-mode (time-lapse imaging) was used to follow the cAMP dynamics overtime, using a 20x, 0.75 N.A. dry objective. Cells were stimulated with 100 nM ISO and with a combination of 25 *μ*M FSK and 100 *μ*M IBMX to maximally raise the cAMP levels. Agonists and inhibitors were added from concentrated stocks and homogeneous distribution was assured through resuspending.

### Image analysis

From Leica LAS X software, the photon arrival times were fitted with a double-exponential reconvolution function using fixed lifetimes of 0.6 ns and 3.4 ns, representing a high-FRET state and low-FRET state, respectively. The generated time-laps images were exported from LAS X as TIF files for ImageJ and further processed and analyzed using a custom macro in Fiji ad detailed before [13]. Cell segmentation was achieved through the deep learning algorithm Cellpose. Single-cell lifetime data were exported as .csv files and analyzed using an R script for visualization and statistical purposes.

### Data presentation and statistical analysis

Data presentation and statistical analysis were conducted using a custom R script developed in RStudio. Graphical representations included a single-cell lifetime graphs, boxplots and bar plots for each sensor to comprehensively illustrate the experimental results. Each trace in the single-cell lifetime graphs represents cAMP dynamics over time for individual cells, offering a detailed visual representation of cellular behavior. To supplement each graph, two boxplots are provided, highlighting the mean values of the traces during the first and last 100 seconds of the experiment. To quantify changes between these two phases, we subtracted the mean values from the first 100 seconds from those of the last 100 seconds per cell, thus considering the standard deviations of the measurements. A Wilcoxon test was conducted to assess the statistical significance of the observed differences. Additionally, the differences in lifetimes were summarized and plotted in a separated barplot, providing a clear comparison of the maximum dynamic range achieved from each sensor, facilitating interpretation of the results.

A binomial statistical test was used to evaluate the tendency for aggregate formation between parental and modified sensors, providing a solid analysis of the observed differences across the experimental conditions.

This approach ensured that the data were presented comprehensively, and that the statistical analysis provided robust conclusions regarding the performance of the new dedicated FLIM sensors in detecting cAMP levels in live cells.

## Supporting information

Supplemental data to Enhanced Epac FLIM sensor

## Acknowledgements

This work has been funded by the European Union’s Horizon Europe research innovation program under Marie Skłodowska-Curie Grant Agreement No. 101073507 (flIMAGIN3D – HORIZON-MSCA-DN-2021, Doctoral Network for a Shared Excellence of Fluorescent Lifetime Imaging Microscopy in Biomedical Applications). We acknowledge the flIMAGIN3D consortium and associated partners for their support. This study has been conducted at The Netherlands Cancer Institute (Amsterdam), which is supported by institutional grants of the Dutch Cancer Society and of the Dutch Ministry of Health, Welfare and Sport. We also thank dr B. vd. Broek and S. Mukherjee in our department for stimulating discussions and for support in automation of data analysis.

