## Supplemental data to Enhanced Epac FLIM sensor for "Cytosolic Enhanced Dark Epac-Based FRET Sensors for Intracellular cAMP detection in live cells via FLIM"

### Supplemental Information

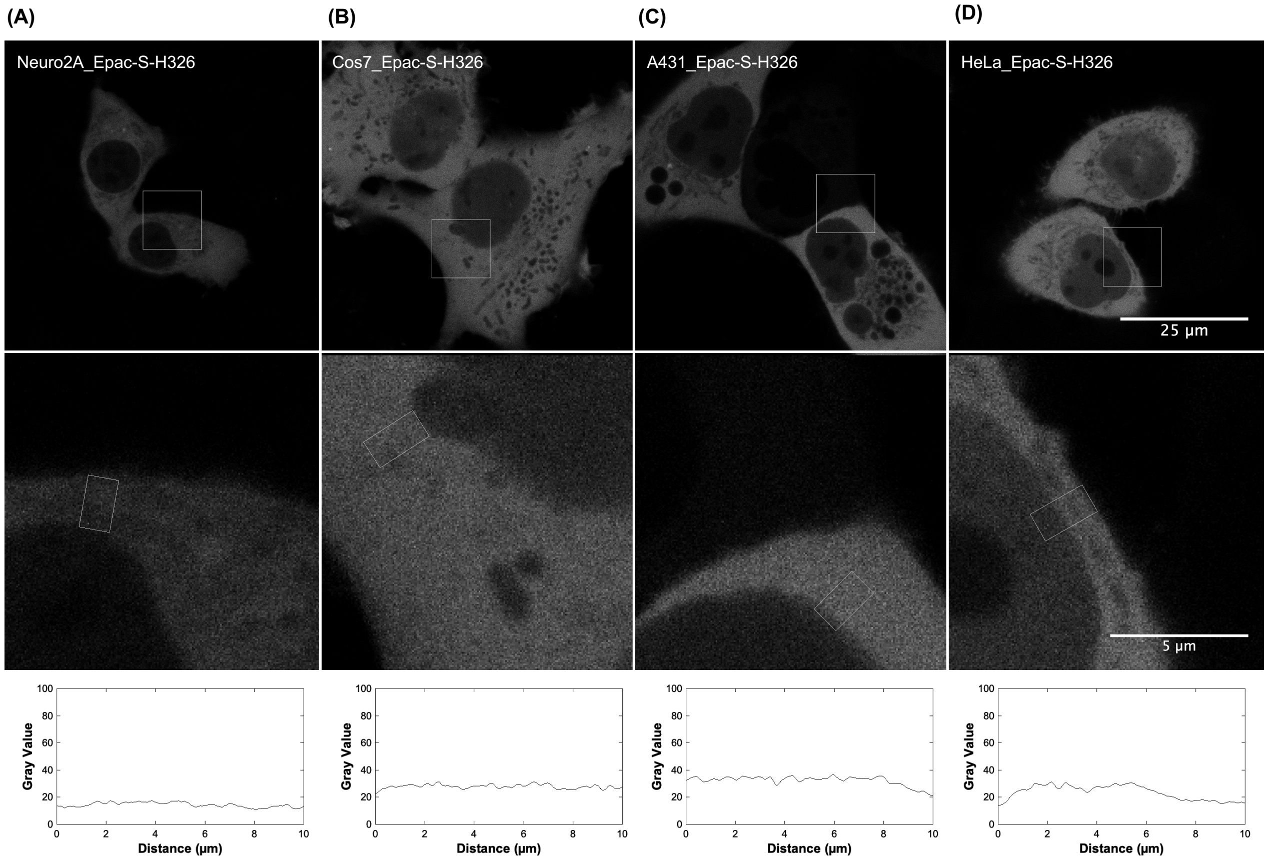


***Fig. S1***

*Fluorescence microscopy images showing the subcellular distribution of Epac-S$^{H326}$ in four distinct cell lines: Neuro2A (**A**), Cos7 (**B**), A431 (**C**), and HeLa (**D**). Magnified views of regions marked by white squares are displayed below each panel. The fluorescence intensity distribution across the selected areas (marked by white rectangles) was quantified using Fiji, and the corresponding distance (μm) versus gray value plots are provided in the bottom panels. Images were captured using a 63x, 1.4 N.A. oil immersion objective.*

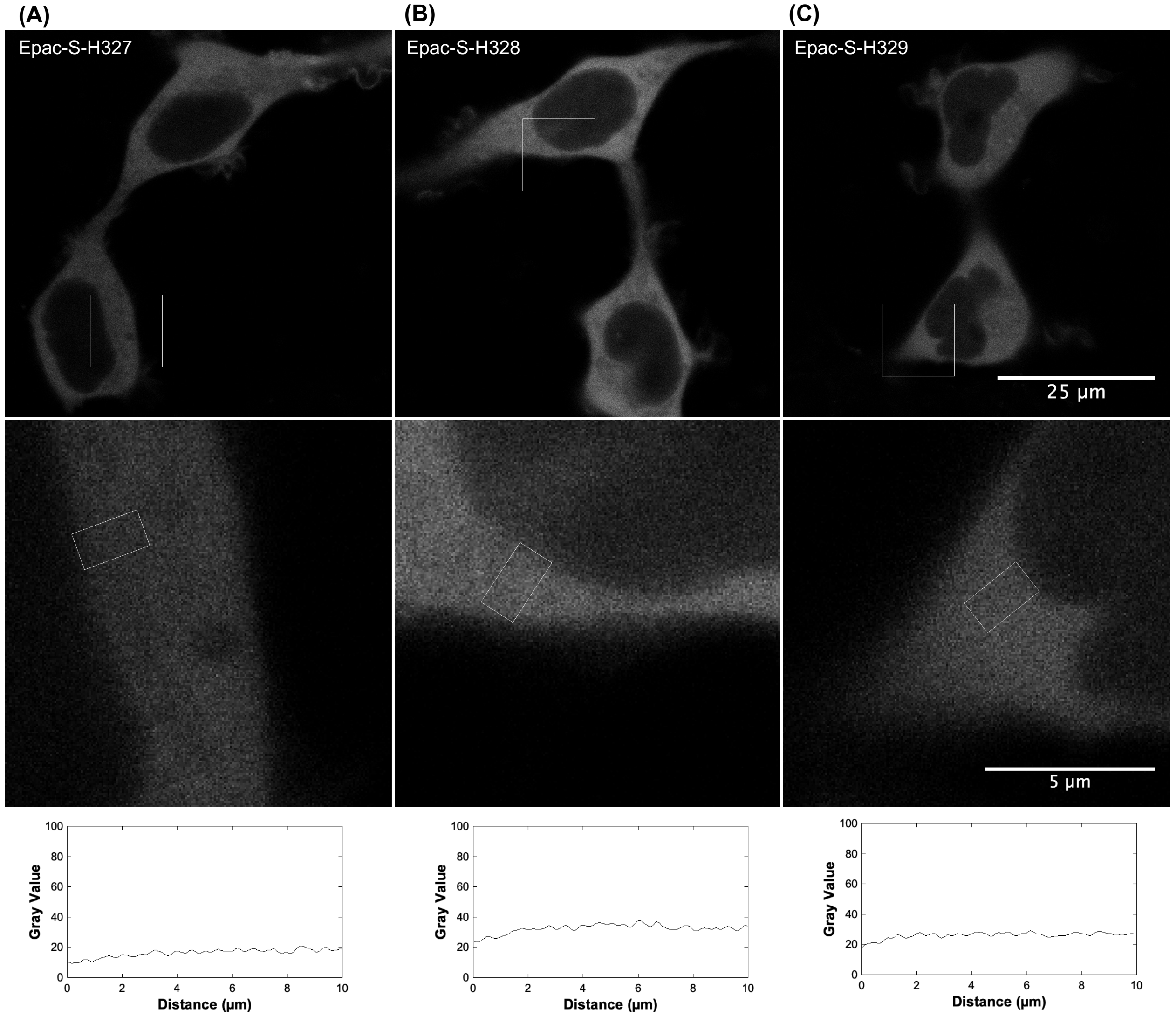


***Fig. S2***

*Microscopy images illustrating the subcellular distribution of Epac sensors within HEK cells. All three dedicated FLIM variants with the highest-affinity, i.e. featuring the M312L (**A**), M312L + E325T (**B**), or E325T (**C**) mutations, exhibit a clear distribution throughout the cytosol without any notable association with the nuclear envelope. The middle row highlights magnified views of the region marked by white square, providing a closer examination of the sensor distribution in these selected area. The bottom panel features plots of distance (μm) versus gray value, derived from intensity profiles within the white rectangular regions, offering a quantitative analysis of the fluorescence distribution across the selected area. 63x, 1.4 N.A. oil immersion objective.*

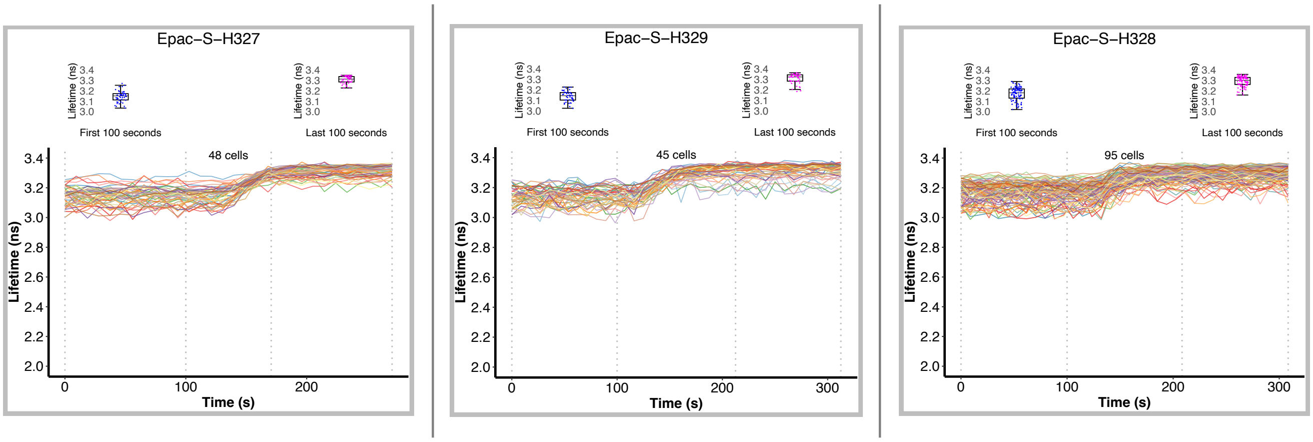


***Fig. S3***

*FLIM analysis of cAMP dynamics in HEK cells expressing high-affinity FRET-FLIM sensors Epac-S$^{H327}$, Epac-S$^{H328}$, Epac-S$^{H329}$. Time-lapse FLIM traces of individual cells, segmented by using the Cellpose algorithm. Cells were monitored at rest and after stimulation with ISO (100 nM) added at 120 seconds and FSK (25 $\mu$M) + IBMX (100 $\mu$M) added at 200 seconds. Due to their high affinity, these sensors are largely saturated at baseline, resulting in a reduced dynamic range. The traces display variations in fluorescence lifetime over time, with corresponding box plots indicating cAMP level changes. 20x, 0.75 N.A. dry objective.*
